# Finger somatotopy is preserved after tetraplegia but deteriorates over time

**DOI:** 10.1101/2021.02.08.430185

**Authors:** Sanne Kikkert, Dario Pfyffer, Michaela Verling, Patrick Freund, Nicole Wenderoth

## Abstract

Following spinal cord injury (SCI), the motor output flow to the limb(s) and sensory input to the brain is largely lost. While attempted movements with the paralysed and sensory deprived body part can still evoke signals in the sensorimotor system, this task-related “net” brain activity of SCI patients differs substantially from healthy controls. Such reorganised and/or altered activity is thought to reflect abnormal processing. It is however possible that this altered ‘net’ sensorimotor activity in SCI patients conceals preserved somatotopically-specific representations of the paralysed and sensory deprived body parts that could be exploited in a functionally meaningful manner (e.g. via neuroprosthetics).

In this cross-sectional study, we investigated whether a functional connection between the periphery and the brain is necessary to maintain somatosensory representations. We used functional MRI and an (attempted) finger movement task to characterise the somatotopic hand layout in the primary somatosensory cortex and structural MRI to assess spared spinal tissue bridges. We tested 14 tetraplegic SCI patients (mean age ± s.e.m.=55 ± 3.6; 1 female) who differed in terms of lesion completeness, retained sensorimotor functioning, and time since injury, as well as 18 healthy control participants (mean age ± s.e.m.=56 ± 3.6 years; 1 female).

Our results revealed somatotopically organised representations of patients’ hands in which neighbouring clusters showed selectivity for neighbouring fingers in contralateral S1, qualitatively similar to those observed in healthy controls. To quantify whether these representations were normal in tetraplegic SCI patients we correlated each participant’s intricate representational distance pattern across all fingers (revealed using representational similarity analysis) with a canonical inter-finger distance pattern obtained from an independent sample. The resulting hand representation typicality scores were not significantly different between patients and controls. This was even true when considering two individual patients with no sensory hand functioning, no hand motor functioning, and no spared spinal tissue bridges. However, a correlational analysis revealed that over years since SCI the hand representation typicality in primary somatosensory cortex deteriorates.

We show that somatosensory representations can be maintained for several years following SCI even in the absence of perhiperhal inputs. Such preserved cortical hand representations could therefore be exploited in a functionally meaningful way by rehabilitation approaches that attempt to establish new functional connections between the hand and the brain after an SCI (e.g. through neuroprosthetics). However, time since injury may critically influence the somatotopic representations of SCI patients and might thereby impact the success of such rehabilitation approaches.

## Introduction

Following a tetraplegic spinal cord injury (SCI; or tetraplegia), individuals mostly experience a loss of muscle function and sensation in their limbs and torso.^1–3^ Accordingly, the primary somatosensory cortex (S1) is mostly deprived of sensory inputs and exposed to altered motor behaviour ^4^. Seminal research in nonhuman primate models of SCI has shown that this leads to extensive cortical reorganisation, in which representations of cortically adjacent body parts (e.g. of the face) take over the deprived brain territory (e.g. of the hand).^5–8^

Human neuroimaging studies have also shown that S1 activity patterns are altered following SCI. In agreement with the non-human primate literature^5–8^, human TMS and neuroimaging suggests that cortically neighbouring body part representations shift towards (but not invade) the deprived M1 and S1 cortex, though results are mixed.^9–14^ Such reorganisation of affected areas in human has been thought to drive both recovery of function and the formation of maladaptive neuronal circuitry that relate to neuropathic pain. While attempted movements with the paralysed and sensory deprived body part are known to still evoke signals in the sensorimotor system, this activity differs substantially from healthy controls^10,13–18^: SCI patients’ volume of activation was found to be reduced^15^, activity levels were increased^13,17,19^, activation was observed in cortical areas that were silent in controls^10,14,15,18,19^, and activity was poorly modulated when task demands changed^15^. This altered sensorimotor activity is attributed to abnormal processing caused by the SCI such as chronic pain^10,12^, compensatory mechanisms^9,13^, overactivation through de-differentiation^14,16^, axonal sprouting of laterally projecting neurons or spinal interneurons^9,20^, or disorganised sensorimotor processing^13,15^. It is possible that this altered ‘net’ sensorimotor processing conceals a preserved and typical somatotopically specific representation of the paralysed and sensory deprived body parts that could be exploited in a functionally meaningful manner (e.g. via neuroprosthetics).

Case studies using intracortical stimulation in the S1 hand area of SCI patients hint at such preserved somatotopic representations, though results are mixed.^21–23^ Negative results were suggested to be due to a loss of hand somatotopy and/or reorganisation in S1 of the implanted SCI patient.^23^ Whether a fine-grained somatotopy of the paralysed body part, which could be exploited in a functionally meaningful manner, is generally preserved in the tetraplegic SCI patient population remains unknown. It is also unclear what clinical, behavioural, and structural determinants may influence such representations to be maintained.

Here we used functional MRI (fMRI) and an attempted finger movement task in tetraplegic SCI patients to examine whether hand somatotopy is preserved following a disconnection between the brain and the periphery. A similar approach has previously been used to map the preserved somatotopic representation of amputees’ missing hands in S1 using volitional phantom finger movements.^24,25^ However, in amputees, these movements typically recruit the residual arm muscles that used to control the missing limb via intact connections between the brain and spinal cord. Whether similar preserved somatotopic mapping can be observed in SCI patients with diminished or no connections between the brain and the periphery is unclear. By measuring a group of tetraplegic SCI patients with varying amounts of spared tissue at the lesion level (quantified by means of midsagittal tissue bridges based on sagittal T2w scans) we uniquely assessed whether preserved connections between the brain and periphery are necessary to preserve fine somatotopic mapping in S1.^26,27^ We also investigated what clinical and behavioural determinants may contribute to preserving S1 somatotopy after chronic SCI.

## Materials and methods

### Participants

Fifteen chronic (i.e. >6 months post injury) tetraplegic SCI patients were recruited and fourteen patients completed the measurements (mean age ± s.e.m.=55 ± 3.6 years; 1 female; 6 dominant left handers; see Table 1 for demographic and clinical details). Eighteen age-, sex-, and handedness-matched able-bodied control participants (age=56 ± 3.6 years; 1 female; 5 dominant left-handers) participated also in this study. Participants’ informed consent was obtained according to the Declaration of Helsinki prior to study onset. Ethical approval was granted by the Kantonale Ethikkommission Zürich (EK-2018-00937). This study is registered on clinicaltrials.gov under NCT03772548. Two patients and one control participant were scanned twice due to excessive head motion during fMRI acquisition or suboptimal slice placement. One patient withdrew from the study prior to study completion. Data of one control participant was distorted and not usable for analysis. All data relating to these subjects was discarded from all data analysis. Patient S02 did not complete the travelling wave measurements due to time constraints.

**Table 1:** Demographic and clinical details. Participants are ordered according to their retained upper limb sensory and motor function (assessed using the Graded Redefined Assessment of Strength, Sensibility and Prehension test; GRASSP). Sex: F=female, M=male; Age=age in years; AIS grade=American Spinal Injury Association (ASIA) Impairment Scale grade defined based on the International Standards for Neurological Classification of Spinal Cord Injury (ISNCSCI), A=complete, B=sensory incomplete, C=motor incomplete, D=motor incomplete, E=normal; Neurological level of injury=defined based on the ISNCSCI; Dominant hand, defined using the Edinburgh handedness inventory: L=left, R=right; GRASSP=Graded Redefined Assessment of Strength, Sensibility and Prehension (maximum score: 232 points); Tested side=side with the lowest score on the GRASSP measurement; GRASSP motor/sensory score of the tested upper limb (maximum scores: 50/24).

|  | Sex | Age | Years<br>since<br>injury | AIS<br>grade | Cause<br>of<br>injury | Neurological<br>level of<br>injury | Dominant<br>hand | GRASSP<br>score | Hand<br>tested | GRASSP<br>tested side<br>motor/sensory |
| --- | --- | --- | --- | --- | --- | --- | --- | --- | --- | --- |
| S01 | M | 32 | 4 | A | trauma | C4 | L | 21 | L | 9/0 |
| S02 | M | 52 | 32 | A | trauma | C5 | R | 78 | L | 16/5 |
| S03 | M | 35 | 4 | A | trauma | C4 | R | 90 | L | 16/17 |
| S04 | M | 41 | 19 | A | trauma | C6 | L | 105 | L | 19/11 |
| S05 | M | 52 | 10 | A | trauma | C2 | L | 118 | R | 23/0 |
| S06 | M | 67 | 26 | A | trauma | C4 | R | 119 | L | 29/2 |
| S07 | M | 57 | 33 | C | trauma | C5 | L | 145 | R | 25/19 |
| S08 | F | 67 | 4 | D | trauma | C5 | L | 173 | L | 41/9 |
| S09 | M | 59 | 12 | D | trauma | C2 | R | 187 | R | 43/9 |
| S10 | M | 42 | 2 | D | trauma | C4 | R | 187 | R | 41/13 |
| S11 | M | 58 | 0.5 | D | ischemic | C4 | R | 194 | R | 37/17 |
| S12 | M | 71 | 16 | D | trauma | C7 | R | 196 | R | 32/24 |
| S13 | M | 65 | 1 | D | trauma | C2 | R | 218 | R | 42/24 |
| S14 | M | 74 | 6 | D | surgery | C4 | L | 220 | R | 47/24 |

### Clinical characterisation

Behavioural testing was conducted in a separate session. We used the International Standards for Neurological Classification of Spinal Cord Injury (ISNCSCI) to neurologically classify patients’ completeness of injury and impairment level. We used the Graded Redefined Assessment of Strength, Sensibility and Prehension (GRASSP) assessment to define sensory and motor integrity of the upper limbs.^28^ Each limb’s maximum score is 116, and refers to healthy conditions.^29^ We determined each patient’s most impaired upper limb according to the GRASSP. Note that GRASSP motor scores reflects overall upper limb motor function (i.e. incl arm and shoulder functioning; see Supplementary Table 1 for muscle specific GRASSP scores). GRASSP sensory scores are hand specific.

### fMRI tasks

We employed two separate paradigms to uncover fine grained somatotopic hand representations using fMRI: First, we used a travelling wave paradigm to investigate the somatotopic hand layout on the S1 cortical surface.^30,31^ Second, we employed a blocked design and representational similarity analysis (RSA) that takes into account the entire fine-grained activity pattern of each finger (i.e. including the representational inter-finger relationships).^32,33^

Participants were visually cued to perform individual finger movements while their palm was positioned up. Patients were instructed to perform the fMRI tasks with their most impaired upper limb (identified using the GRASSP, see above). Controls’ tested hands were matched to the patients. Due to their injury, not all patients were able to make overt finger movements. In these cases, patients were carefully instructed by the experimenter to make attempted (i.e. not imagined) finger movements. The experimenter explained that although it is not possible for the patient to perform an overt movement, attempting to perform the movement may still produce a motor command in the brain. Importantly, complete paraplegic SCI patients are able to distinguish between attempted and imagined movements with their paralysed body part.^15,18,34^ Furthermore, attempted movements activated SCI patients’ motor network similarly to controls performing overt foot movements.^15,18,34^

Participants saw five horizontally aligned white circles, corresponding to the five fingers, via a visual display viewed through a mirror mounted on the head coil. For participants moving their left hand, the leftmost and rightmost circles corresponded to the thumb and little finger, respectively. For participants moving their right hand, the leftmost circle corresponded to the little finger and the rightmost circle to the thumb. To cue a finger movement, the circle corresponding to this finger turned red. Participants performed self-paced flexion/extension movements with the cued finger for the duration of the colour change. Instructions were delivered using Psychtoolbox (v3) implemented in Matlab (v2014). Head motion was minimized using over-ear MRI-safe headphones or padded cushions.

The travelling wave paradigm involved individuated finger movements in a set sequence. Each 10s finger movement block was immediately followed by a movement block of a neighbouring finger. The forward sequence cycled through the fingers: thumb-index-middle-ring-little. To account for order-related biases due to the set movement cycle and sluggish hemodynamic response, we also collected data using a backward sequence: The backward sequence cycled through the movements in a reverse of the forward sequence: little-ring-middle-index-thumb fingers. The forward and backward sequences were employed in separate runs. A run lasted 6min and 4s, during which a sequence was repeated seven times. The forward and backward runs were repeated twice, with a total duration of 24min and 16s.

The blocked design consisted of six conditions: Movement conditions for each of the five fingers and a rest condition. Finger movement instructions were as described above and the word “Rest” indicated the rest condition. A block lasted 8s and each condition was repeated five times per run in a counterbalanced order. Each run comprised a different block order and had a duration of 4min and 14s. We acquired four runs, with a total duration of 16min and 56s.

### MRI acquisition

MRI data was acquired using a Philips 3 tesla Ingenia system (Best, The Netherlands) with a 20-channel HeadNeckSpine or, in case of participant discomfort due to the coil’s narrowness, a 15-channel HeadSpine coil. Anatomical T1-weighted images covering the brain and cervical spinal cord were acquired using the following acquisition parameters: 0.8mm^3^ resolution, repetition time (TR): 9.3ms, echo time (TE): 4.4ms, flip angle: 8°. Anatomical T2-weighted images of the cervical spinal cord were acquired sagitally using the following acquisition parameters: 1×1×3mm resolution, TR: 4500ms, TE: 85ms, flip angle: 90°, slice gap: 0.3mm, 15 slices. Task-fMRI data was acquired using an echo-planar-imaging (EPI) sequence with partial brain coverage. 22 sagittal slices were centred on the anatomical location of the hand knob with coverage over the thalamus and brainstem. We used the following parameters: 2mm^3^ resolution, TR: 2000ms, TE: 30ms, flip angle: 82°, SENSE factor: 2.2. We acquired 182 and 127 volumes for each of the travelling wave and blocked design runs, respectively.

### fMRI analysis

fMRI analysis was implemented using FSL v6.0 (https://fsl.fmrib.ox.ac.uk/fsl/fslwiki) Advanced Normalization Tools (ANTs) v2.3.1 (http://stnava.github.io/ANTs/), the RSA toolbox^35,36^, and Matlab (R2018a). Cortical surface visualisations were realised using Freesurfer (https://surfer.nmr.mgh.harvard.edu/)^37,38^ and Connectome Workbench (https://www.humanconnectome.org/software/connectome-workbench).

#### fMRI preprocessing

Common preprocessing steps were applied using FSL’s Expert Analysis Tool (FEAT). The following preprocessing steps were included: motion correction using MCFLIRT^39^, brain extraction using automated brain extraction tool BET^40^, spatial smoothing using a 2mm full-width-at-half-maximum (FWHM) Gaussian kernel, and 100s high-pass temporal filtering with a 100s (blocked design runs) or 90s (traveling wave runs) cut-off.

#### Image registration

Image coregistration was done in separate, visually inspected, steps. For each participant, a midspace was calculated between the four blocked design runs, i.e. an average space in which images are minimally reoriented. We then transformed all fMRI data to this midspace using purely rigid probability mapping in ANTs. Next, we registered each participant’s midspace to the T1-weighted image, initially using 6 degrees of freedom and the mutual information cost function, and then optimised using boundary based registration (BBR).^41^ Each coregistration step was visually inspected and, if needed, manually optimised using blink comparison in Freeview.

#### Travelling wave analysis

The travelling wave approach is characterised by set finger movement cycles that are expected to result in neighbouring cortical activations. It is designed to capture voxels that show preferential activity to one condition, above and beyond all other conditions (i.e. winner-takes-all principle; finger selectivity). The travelling wave approach is especially powerful to reveal the smooth progression of neighbouring representations that are specific for topographic maps. This technique is therefore frequently used to uncover retinotopic^42^, somatotopic^25,30,31,43,44^, and tonotopic representations^45,46^. Importantly, S1 finger movement somatopy is highly consistency across multiple travelling wave scanning sessions.^31^

Travelling wave analysis was conducted separately for each participant, closely following procedures previously described in Kikkert et al.^25^ A reference model was created using a gamma hemodynamic response function (HRF) convolved boxcar, using a 10s ‘on’ (a single finger movement duration) and 40s ‘off’ period (movement duration of all other fingers). This reference model was then shifted in time to model activity throughout the full movement cycle. Since we had a 2s TR and a 50s movement cycle, the reference model was shifted 25 times.

Within each individual run, each voxel’s preprocessed BOLD signal time-course was cross-correlated with each of the 25 reference models. This resulted in 25 r-values per voxel per run that were normalised using a Fisher r-to-z transformation. To create finger-specific (i.e. hard-edged) maps, we assigned individual lags to specific fingers and averaged the r-values across these lags. This resulted in five r-values, one for each finger, per voxel per run. These r-values were averaged across runs per voxel and finger assignment. As a result, each individual voxel now contained an averaged r-value for each finger. Next, a winner-take-all approach to assign each voxel to one finger exclusively based on the maximum r-value, providing us with finger-specificity.

To visualise the smooth gradient of progression across fingers we produced lag-specific maps. The backward run’s standardised cross-correlation r-values were lag-reversed and averaged with the forward runs per voxel and per lag. This resulted in 25 r-values per voxel (one per lag). Next, we used a winner-take-all principle to find the maximum r-value across lags for each voxel, providing us with lag-specificity.

Cortical surface projections were constructed from participant’s T1-weighted images. The finger-specific maps and lag-specific gradient maps were projected onto the cortical surface using cortical-ribbon mapping. Thresholding was applied on the cortical surface using a false discovery criterion q < 0.05 based on the native (3D) values. The FDR-thresholded finger-specific maps were combined to form a hand map. Within this hand map, the lag-specific map was used to visualise the smooth gradient of progression across fingers. We were unable to find a characteristic hand map in one patient who reported post hoc that he performed imagined movements during the measurement.

#### Spatial correspondence of finger maps over time: Dice overlap coefficient analysis

To confirm that the travelling wave finger-specific maps did not represent random noise we quantified spatial consistency of finger preference between two halves of the data using the Dice overlap coefficient (DOC).^25,47,48^ The DOC calculates the spatial overlap between two representations relative to the total area of these representations. The DOC ranges from 0 (no spatial overlap) to 1 (perfect spatial overlap). If A and B represent the areas of two representations, then the DOC is expressed as:

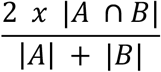

The averaged finger-specific maps of the first forward and backward runs formed the first data half. The averaged finger-specific maps of the second forward and backward runs formed the second data half. The finger-specific clusters were minimally thresholded (Z>2) on the cortical surface and masked using an S1 ROI, created based on Brodmann area parcellation using Freesurfer. The DOC was calculated between each possible finger pair across the two data halves (see Fig. 1C). A square root transformation was applied to the resulting non-normally distributed DOC scores.

**Figure 1:**
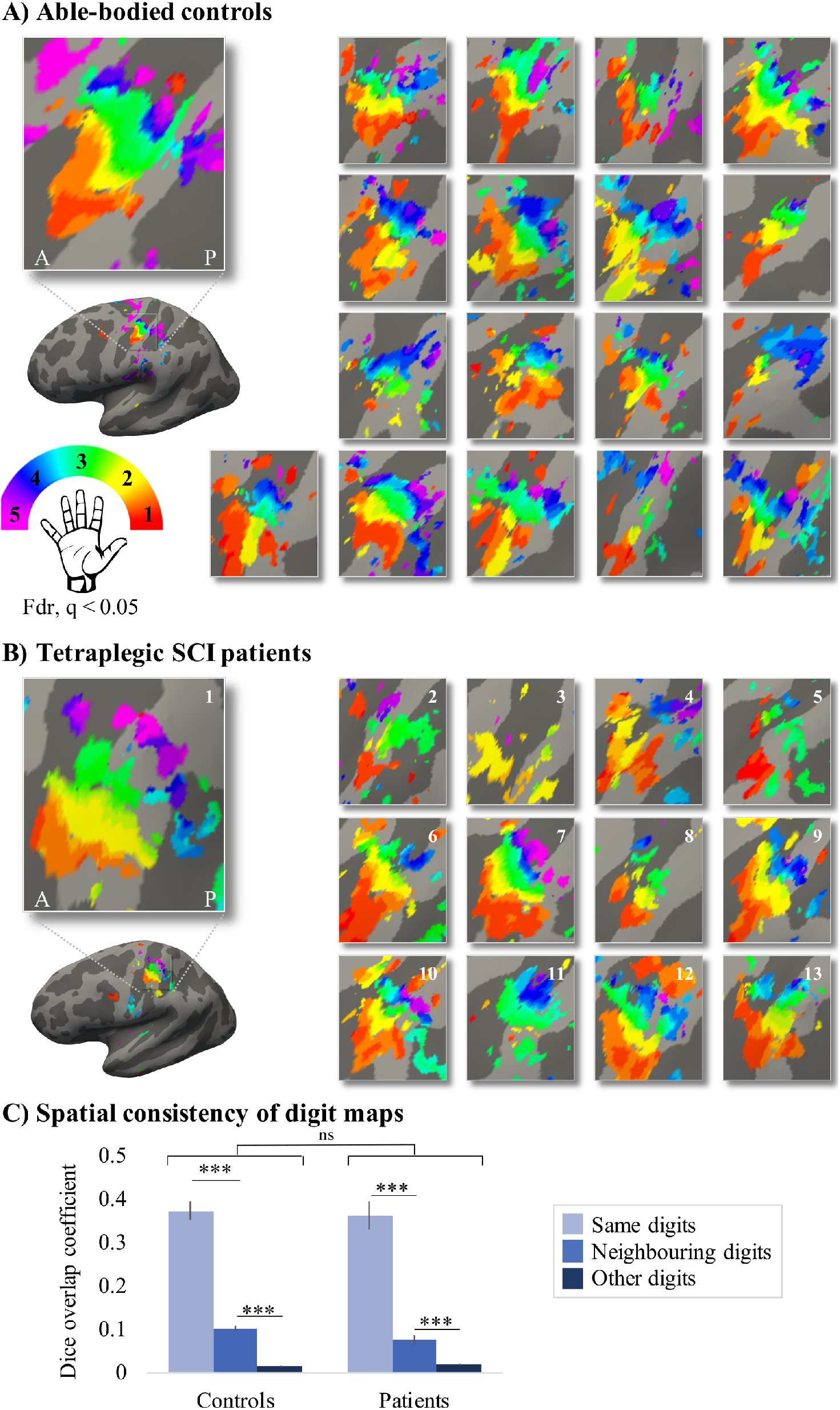
Finger selectivity is preserved in tetraplegic SCI patients. Colours indicate selectivity for the thumb (digit 1, red), index finger (digit 2, yellow), middle finger (digit 3, green), ring finger (digit 4, blue), and little finger (digit 5, purple). Typical finger selectivity is characterized by a gradient of finger preference, progressing from the thumb (laterally) to the little finger (medially). These typical digit gradients can be observed in both the able-bodied controls (A) and the tetraplegic SCI patients (B). Qualitatively, the position, finger order, and extent of the patient maps were generally similar to those observed in controls. Patients’ hand maps are sorted according to their upper-limb impairments (assessed using the GRASSP): from most to least impaired - as indicated by the white numbers. White arrows indicate the central sulcus. A=anterior; P=posterior. Multiple comparisons were adjusted using a false discovery rate (FDR) with q < 0.05. C) To ensure that the observed clusters were not representing noise, but rather true finger selectivity, we calculated split-half consistency between two halves of the travelling wave dataset. Both controls and patients showed higher split-half consistency (assessed using the Dice overlap coefficient) for comparison of the same fingers between two halves of the travelling wave dataset (light blue), compared to neighbouring (blue) and other fingers (dark blue). Moreover, neighbouring fingers showed greater overlap across the split-halves of the dataset then non-neighbouring fingers for both patients and controls. Error bars show the standard error of the mean. Significance is indicated by: ***=corrected p ≤ 0.001, ns=non-significant.

If the finger maps would be spatially consistent and represent true finger selectivity, then one would expect a higher DOC between pairs of ‘same’ fingers across the two data halves compared to neighbouring and non-neighbouring finger pairs. One would further expect to find a somatotopic relationship in the DOCs: i.e. a higher DOC between neighbouring compared to non-neighbouring finger pairs.

#### Univariate analyisis

To assess univariate task-related activity of the blocked design data, time-series statistical analysis was carried out per run using FMRIB’s Improved Linear Model (FILM) with local autocorrelation correction, as implemented in FEAT. We obtained activity estimates using a general linear modelling (GLM) based on the double-gamma HRF and its temporal derivative. Each finger movement conditions was contrasted with rest. A further contrast was defined for overall task-related activity by contrasting all movement conditions with rest.

We defined an S1 hand ROI by converting the split-half consistency S1 ROI to volumetric space. Any holes were filled and non-zero voxels were mean dilated. Next, the axial slices spanning 2cm medial/lateral to the hand knob^49^ were identified on the 2mm MNI standard brain (min-max MNI z-coordinates=40-62). This mask was non-linearly transformed to each participant’s native structural space. Finally, we used this mask to restrict the S1 ROI and extracted an S1 hand area ROI. The average BOLD response for overall task-related activity was extracted per run for voxels underlying this S1 hand ROI and averaged across runs per participant. A similar analysis was used to investigate overall task-related activity in an M1 hand ROI (see Supplementary Fig. 1).

#### Representational similarity analysis

While the traveling wave approach is powerful to uncover the somatotopic finger arrangement, a fuller description of hand representation can be obtained by taking into account the entire fine-grained activity pattern of all fingers. We therefore computed the dissimilarity between the activity patterns measured for each finger pair within the S1 hand ROI using the cross-validated squared Mahalanobis distance (or crossnobis distance).^35^ We extracted the blocked design voxel-wise parameter estimates (betas) for each condition versus rest (identified in the univariate analysis) and the model fit residuals under the S1 hand ROI. We prewhitened the betas using the model fit residuals. We then calculated the cross-validated squared Mahalanobis distances between each possible finger pair, using our 4 runs as independent crossvalidation folds, and averaged the resulting distances across the folds. If it is impossible to statistically differentiate between conditions (i.e. when this parameter is not represented in the ROI), the expected value of the distance estimate would be 0. If it is possible to distinguish between activity patterns this value will be larger than 0.

The dissimilarity values for all finger pairs were assembled in a representational dissimilarity matrix (RDM), with a width and height corresponding to the 5 finger movement conditions. All statistical analysis was conducted on the unique values of the RDM. We estimated the strength of the finger representation or “finger separability” by averaging the 10 unique off-diagonal values of the RDM. If there is no finger information in the ROI, the finger separability would be 0. Second, we estimated the typicality (or normality) of each participant’s RDM by calculating a Spearman correlation with a canonical RDM. This canonical RDM was based on 7T finger movement fMRI data in an independently acquired cohort of healthy controls (n=8). The S1 hand ROI used to calculated this canonical RDM was defined similarly as in the current study (see Wesselink and Maimon-Mor^50^ for details). The typicality scores were Fisher r-to-z transformed prior to statistical analysis (the r*s* typicality scores are used solely for visualisation). Controls’ and SCI patients’ typicality scores were compared to those of a group of individuals with congenital hand malformation (n=13), hereafter one-handers, obtained in another study.^24^ Congenital one-handers are born without an arm and do not have an S1 hand representation contralateral to the missing hand. Importantly, these typicality scores were calculated using the same procedures described above.

Finally, we performed multidimensional scaling (MDS) to visualise the dissimilarity structure of the RDM in an intuitive manner. MDS projects the higher-dimensional RDM into a lower-dimensional space, while preserving the inter-finger dissimilarity values as well as possible.^51^ MDS was performed for each individual participant and then averaged per group after Procrustes alignment to remove arbitrary rotation induced by MDS.

### Structural MRI analysis

#### Midsaggtical tissue bridges analysis

We used sagittal T2w structural images of the cervical spinal cord at the lesion level to quantify spared tissue bridges. It has been shown that, already in the sub-acute stage after SCI, edema and haemorrhage have largely resolved and hyperintense signal changes reliably reflect intramedullary neural damage.^26,27,52^ We closely followed previously described procedures.^26,27,52^ We used Jim 7.0 software (Xinapse Systems, Aldwincle, UK) for manual lesion segmentation at the lesion level, for which high intra- and interobserver reliability has previously been reported.^26,27^ The experimenter conducting the manual segmentation was blinded to patient identity. We only included patients’ T2w scans if the lesion (i.e. hyperintense, cerebrospinal fluid filled cystic cavity) was clearly visible on the midsagittal slice. We excluded images with metal artefacts or insufficient quality which would not allow a reliable quantification of lesion measures. Tissue bridges were defined as the relatively hypointense intramedullary region between the hyperintense CSF on one side and the cystic cavity on the other side. We assessed the width of ventral and dorsal tissue bridges on the midsagittal slice and summed these to get the total width of tissue bridges.

#### Cervical cross-sectional spinal cord area analysis

We used Jim 7.0 software (Xinapse Systems, Aldwincle, UK) to extract the cross-sectional spinal cord area at cervical level C2/3 of the spinal cord from the sagittal T2w scans. We used multi-planar reconstruction of sagittal images resulting in 10 contiguous axial slices at C2/3 with a thickness of 2 mm.^53^ Using the active-surface model from Horsfield et al.^54^, the cross-sectional spinal cord area was calculated semi-automatically for every slice and averaged over all 10 slices.

### Statistical analysis

Statistical analysis was carried out using SPSS (v25) Standard approaches were used for statistical analysis, as mentioned in the Results section. If normality was violated (assessed using the Shapiro-Wilk test), non-parametric statistical testing was used. We used a Crawford-Howell t-test to compare single patients to the congenital and control groups.^55^ All testing was two-tailed and corrected p-values were calculated using the Benjamini-Hochberg procedure to control the FDR with q < 0.05. The correlational analysis was considered exploratory and we did not correct for multiple comparisons in this analysis.

Bayesian analysis was carried out using JASP (v0.12.2) for the main comparisons to investigate support for the null hypothesis with the Cauchy prior width set at 0.707 (i.e. JASP’s default). Following the conventional cut-offs, a Bayes Factor (BF) smaller than 1/3 is considered substantial evidence in favour of the null hypothesis. A BF greater than 3 is considered substantial evidence and a BF greater than 10 is considered strong evidence in favour of the alternative hypothesis. A BF between 1/3 and 3 is considered weak or anecdotal evidence.^56,57^

## Data availability

Full details of the experimental protocol are available on clinicaltrials.gov under the number NCT03772548. Data is shared on <*link will be included upon publication*>.

## Results

### Patient impairments

We tested a heterogenous group of SCI patients in terms of completeness of the SCI (ranging from AIS-A to AIS-D), neurological level of the injury (ranging from C2 to C7), years since injury (ranging from 6 months to 33 years since SCI), and sensorimotor impairments (ranging from a GRASSP score of 21 to 220; healthy GRASSP score=232).

### Finger selectivity is preserved following tetraplegic spinal cord injury

Using a travelling wave approach, we found detailed somatotopy maps of SCI patients’ hands, in which neighbouring clusters showed selectivity for neighbouring fingers in contralateral S1, similar to those observed in healthy controls (see Fig. 1 A&B). A characteristic hand map shows a gradient of finger preference, progressing from the thumb (red, laterally) to the little finger (pink, medially). Such a characteristic hand map was found even in a patient who suffered complete paralysis and sensory deprivation of the tested hand. Generally, the position, order of finger preference, and extent of the hand maps were qualitatively similar between patients and controls.

To ensure that the observed clusters were not representing noise, but rather true finger selectivity, we calculated split-half consistency between two halves of the dataset using the Dice overlap coefficient. Minimally thresholded finger maps were compared across the split-halves of the data within an S1 mask. Overall, split-half consistency was the same between patients and controls, as tested using a repeated measures ANOVA (see Fig. 1C, note that the data in the Figure is not transformed to easy interpretability; F_(1)_=1.15, p=0.29; BF_10_=0.27). There was a difference in split-half consistency between pairs of same, neighbouring, and non-neighbouring fingers (F_(1.52)_=410.32, p < 0.001; BF_10_=5.82e +43). This neighbourhood relationship was not significantly different between the control and patient groups (i.e. there was no significant interaction; F_(1.52)_=1.85, p=0.18; BF_10_=0.67), with the Bayes Factor (BF) showing anecdotal evidence in favour of the null hypothesis. The Dice overlap coefficient was highest for comparison of the same fingers between two halves of the dataset compared to neighbouring (controls: t_(17)_=15.51, p < 0.001; patients: t_(12)_=15.36, p < 0.001), and non-neighbouring fingers (controls: t_(17)_=23.31, p < 0.001; patients: t_(12)_=11.45, p < 0.001). Moreover, neighbouring fingers showed greater overlap across the split-halves of the dataset than non-neighbouring fingers (controls: t_(17)_=16.57, p < 0.001; patients: t_(12)_=4.57, p=0.001). This demonstrates that there was a somatotopic gradient in split-half consistency that was similar between the control and patient groups.

### Typical hand somatotopy is preserved following tetraplegic spinal cord injury

Next, we assessed univariate task-related activity of the blocked design data, quantified by averaging the finger movement BOLD responses versus baseline across all fingers within the contralateral S1 hand ROI (see Fig. 2A). Overall, all patients were able to engage their S1 hand area by moving individual fingers (t_(13)_=9.16, p < 0.001; BF_10_=3.26e +4), as did controls (t_(17)_=9.89, p < 0.001; BF_10_=7.05e +5). Furthermore, patients’ task-related activity was not significantly different from controls (t_(30)_=−0.35, p=0.73; BF_10_=0.36), with the BF showing anecdotal evidence in favour of the null hypothesis. Similar results were found when exploring univariate task-related activity in the contralateral M1 hand ROI (see Supplementary Fig. 1).

**Figure 2:**
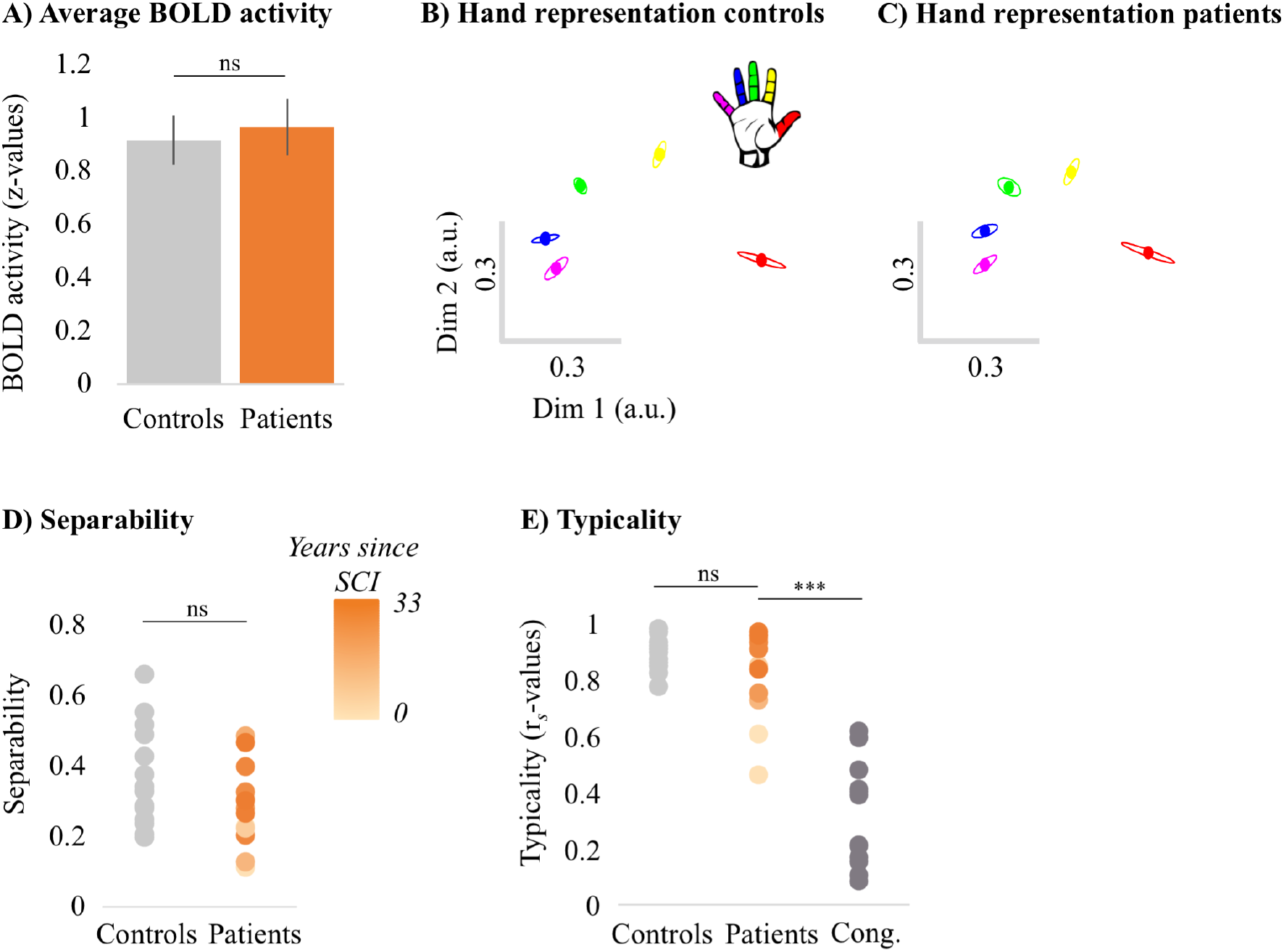
Typical multivariate hand somatotopy is preserved following tetraplegic spinal cord injury. A) Task-related activity in the S1 hand ROI for able-bodied controls (grey) and tetraplegic SCI patients (orange). B-C) Two-dimensional projection of the representational structure of the hand in the control (B) and patient groups (C). Dissimilarity is reflected by the distance in the two dimensions. Individual digits are represented by different colours: thumb=red; index finger=yellow; middle finger=green; ring finger=blue; little finger=purple. Ellipses represent the between-subject standard error after Procrustes alignment. D) Separability, measured as mean dissimilarity, of the representational structure in the S1 hand area of controls and patients. Patients are presented on a colour scale representing the sensory and motor functioning of their tested upper limb, measured using the GRASSP test (0=no upper limb function, 116=normal upper limb function). E) Typicality of the representational structure in controls, patients, and congenital one-handers (Cong. in the figure). Significance is indicated by: ***=p < 0.001; ns=non-significant; Dim=dimension; a.u.= arbitrary unit; Cong=congenital one-handers.

While the travelling wave maps demonstrate finger selectivity, they provide little information about the overlap between finger representations. We examined the intricate relationship between finger representations for all patients and controls using representational similarity analysis (see Figures 2B-C). The resulting inter-finger dissimilarity values were averaged across finger pairs within each participant to obtain an estimate for average inter-finger separability (see Fig. 2D). We found that the inter-finger separability was greater than 0 for patients (t_(13)_=9.83, p < 0.001; BF_10_=6.77e +4) and controls (t_(17)_=11.70, p < 0.001; BF_10_=6.92e +6), indicating that the S1 hand area in both groups contained information about individuated finger representations. We did not find a significant group difference (t_(30)_=1.52, p=0.14; BF_10_=0.81), with the BF showing anecdotal evidence in favour of the null hypothesis.

Although inter-finger separability was not significantly different between patients and controls, it is possible that the pattern of individuated finger activity was atypical in the patients. We therefore examined whether the inter-finger distance pattern was normal in tetraplegic SCI patients (see Fig. 2E) by correlating each participant’s inter-finger distance pattern with a canonical inter-finger distance pattern. SCI patients’ typicality scores were compared to those of the controls and of a group congenital one-handers (data taken from an independent study^24^). Congenital one-handers are born without a hand and do not have a missing hand representation. This group was therefore included as a control for absence of hand representation. We found a significant difference in typicality between SCI patients, healthy controls, and congenital one-handers (H_(2)_=26.64, p < 0.001). As expected, posthoc tests revealed significantly higher typicality in controls compared to congenital one-handers (U=0, p < 0.001; BF_10_=113.60). Importantly, inter-finger pattern typicality of the SCI patients was significantly higher than the congenital one-handers (U=4, p < 0.001; BF_10_=90.33), but not significantly different from the controls (U=103, p=0.40; BF_10_=0.55). The BF for the comparison between SCI patients and controls showed anecdotal evidence for equivalence between both groups.

### Typical hand somatotopy deteriorates over years after spinal cord injury

Next, we aimed to understand which clinical, behavioural, and structural determinants may allow hand representations in S1 to be maintained. We first explored correlations with S1 hand representation typicality. We found that the number of years since SCI significantly correlated with hand representation typicality (see Fig. 3A; r_*s*_=−0.59, p=0.028), suggesting that S1 hand representation typicality may deteriorate over time after SCI. We further found patients with more retained GRASSP motor function of the tested upper limb had more typical hand representations in S1 (see Fig. 3B; r_*s*_=0.60, p=0.02). We did not find a significant correlation between S1 hand representation typicality and GRASSP sensory function of the tested upper limb, spared midsagittal tissue bridges, or cross-sectional spinal cord area (see Figures 3C-E; r_*s*_=0.40, p=0.16, r_*s*_=0.46, p=0.14, and r_*s*_=0.33, p=0.25, respectively).

**Figure 3:**
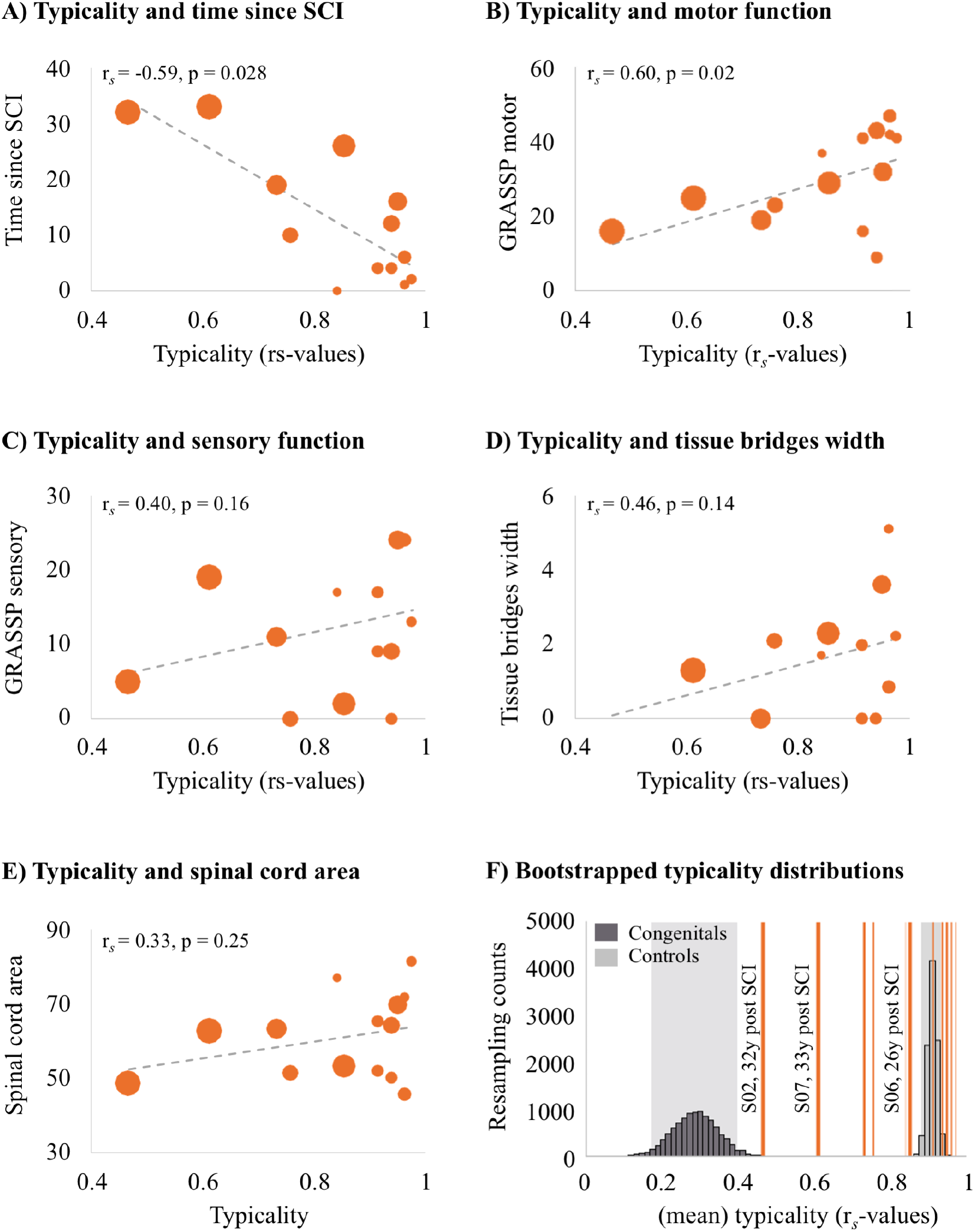
Years since spinal cord injury and retained motor function correlates with hand representation typicality in the primary somatosensory cortex (S1). We examined clinical, behavioural, and structural correlates for hand representation typicality. Increasing marker sizes represent increasing years since SCI in graphs A-E. A) There was a negative correlation between years since SCI and hand representation typicality. B) We found a positive correlation between motor function (measured using the GRASSP) and hand representation typicality. C) There was no significant correlation between sensory function (measured using the GRASSP) and hand representation typicality. There was no significant correlation between hand representation typicality and spared midsagittal tissue bridges (D) or cross-sectional spinal cord area (E). F) Bootstrapped distribution of controls’ and congenital one-handers’ mean S1 hand representation typicality. Dark grey bars indicate the distribution of congenital one-handers (data taken from an independent study^24^), and light grey bars indicate the distribution of the able-bodied controls (tested for this study). The typicality scores of the SCI patients are plotted as orange lines. Increasing line thickness represent increasing years since SCI. Grey shaded areas indicate the 95% confidence intervals of the mean for congenital one-handers and able-bodied controls.

We further explored the hand representation typicality of patients S01 and S03 who did not have any spared midsagittal tissue bridges at the lesion level, a complete (S01) or near complete (S03) hand paralysis, and a complete (S01) or near complete loss (S03) of hand sensory function (as assessed using the GRASSP test). Interestingly, both patients had a highly typical hand representation that was significantly different from congenital one-handers (i.e. who are born without a hand and do not have a missing hand representation; S01: t_(12)=_3.20, p=0.008; S03: t_(12)_ =2.97, p=0.01), but not controls (S01: t_(17)=_0.95, p=0.36; S03: t_(17)_ =0.04, p=0.97). This suggests that retained connections between the periphery and the brain, retained motor functioning, and retained sensory functioning may not be necessary to maintain hand representation typicality.

We then ran an exploratory stepwise linear regression to investigate which of these clinical, behavioural, and structural characteristics were predictive of hand representation typicality in S1. Years since SCI significantly predicted hand representation typicality in S1 with R^2^=0.40 (F_(1,10)_=6.73, p=0.027). Motor function, sensory function, tissue bridges, and spinal cord area did not significantly add to the prediction (t=1.43, p=0.19, t=1.44, p=0.18, t=1.19, p=0.26, and t=0.41, p=0.69, respectively). This analysis suggests that while hand representations are preserved following SCI, they may deteriorate over time.

To inspect this further, we bootstrapped the mean typicality of the able-bodied controls and congenital one-handers 10,000 times to infer the population means. While most SCI patients’ typicality scores fell in or very close to the able-bodied controls’ distribution, we found that some SCI patients’ typicality scores fell in-between the able-bodied controls’ and congenital one-handers’ distributions. This suggests that these patients may not be similar to congenital one-handers or to able-bodied controls. Interestingly, this included those patients for whom most years had passed since their SCI. This suggests that S1 hand representations might deteriorate after an SCI, but some weak hand information may be maintained in S1 even >30 years after an SCI.

## Discussion

In this study we investigated whether hand somatotopy is preserved following a tetraplegic SCI. We tested a heterogenous group of SCI patients to examine what clinical, behavioural, and structural determinants contribute to preserving S1 somatotopy. Our results revealed detailed somatotopically organised finger maps of tetraplegic SCI patients’ hands in which neighbouring clusters showed selectivity for neighbouring fingers in contralateral S1, similar to those observed in controls. Correspondingly, we found that the inter-finger relationship patterns in SCI patients were more typical than those of congenital one-handers (i.e. individuals who are born without a hand and do not have a hand representation^24^), but not different from controls.

Crucially, spared spinal midsaggital tissue bridges, motor function, and sensory function did not seem necessary to maintain and activate a somatotopic hand representation in S1. Firstly, these behavioural and structural determinants were not predictive of hand representation typicality. Secondly, we found a highly typical hand representation in two patients (S01 and S03) who did not have any spared spinal tissue bridges at the lesion level, a complete (S01) or near complete (S03) hand paralysis, and a complete (S01) or near complete loss (S03) of hand sensory function. These results are in line with the notion that somatotopic activity patterns in S1 can be elicited through efference copies from the motor system. While motor and sensory signals no longer pass through the spinal cord in the absence of spinal tissue bridges, S1 and M1 remain intact. When a motor command is initiated (e.g. in the form of an attempted hand movement) an efference copy is thought to be sent to S1 in the form of corollary discharge. This corollary discharge is thought to resemble the expected somatosensory feedback activity pattern and may drive somatotopic S1 activity even in the absence of ascending afferent signals from the hand.^58,59^

Time since injury was predictive of a deteriorated, or less typical, somatotopic S1 hand representation. In fact, both patients with a typical S1 hand representations in the absence of spinal tissue bridges suffered their SCI only 4 years ago (the group was on average 12 years since SCI). The hand representation typicality of patients who suffered their SCI further in the past were not similar to congenital one-handers’ or to controls’ hand representation. Thus, S1 hand representations may deteriorate over time after an SCI, but some weak hand information appears to be maintained in S1 even >30 years after an SCI. This finding complements previous studies in amputees showing that missing hand somatotopy is preserved even decades after arm amputation and that years since injury was not related to missing hand representation typicality.^24,25^ While both amputees and SCI patients suffer from major sensory input loss and changed motor behaviour, their injuries are inherently different. The injured axons within amputees’ residual limb and the remaining part of the peripheral nerves mostly generate some spontaneous (ectopic) activity that is propagated to the brain and could contribute to maintaining S1 representations.^25,60–64^ Furthermore, amputees mostly remain able to move and receive afferents from the residual limb muscles that used to control the missing hand, as most of their motor system remains intact. Although their hand is missing amputees’ residual arm muscles are often still used (either to move the residual limb control and/or a prosthetic arm). Lastly, amputees’ vividness of kinaesthetic sensations during phantom finger movements was found to be predictive of the typicality of the S1 missing hand representation.^24^ A continued, though altered, experience relating to the missing hand in amputees may contribute to maintaining the somatotopic missing hand S1 representation. Contrarily, SCI patients mostly have reduced or a complete loss of communication between the brain and periphery. They therefore have problems activating the adequate muscles and will lose orderly afferents from their muscles and the skin. It is possible that this continued disuse causes cortical somatotopic S1 representations of SCI patients to deteriorate after the injury.

How may these representations be preserved over time in the absence of peripheral information? Firstly, it is possible that these somatotopic maps are relatively hardwired and while they deteriorate over time, they never fully disappear. Indeed, somatotopic mapping of a sensory deprived body part has been shown to be resilient after dystonia^65^ (though see^66,67^) and arm amputation^24,25,68^. Second, it is possible that cortico-cortical efference copies may keep a representation ‘alive’ through occasional corollary discharge.^58^ It is possible that our patients occasionally performed attempted movements which would result in corollary discharge in S1. Third, it is possible that even though a patient is clinically assessed to be complete and is unable to perceive sensory stimuli on the deprived body part, there is still some ascending information flow that contributes to preserving somatotopy.^11^ A recent study found that although complete paraplegic SCI patients were unable to perceive a brushing stimulus on their toe, 48% of patients activated the location appropriate S1 area.^11^ However, the authors of this study defined the completeness of patients’ injuries via behavioural testing, while we assessed the retained connections passing through the SCI directly via quantification of spared tissue bridges through structural MRI. It is unlikely that spinal tissue carrying somatotopically organised information would be missed by our assessment.^26,27^ Fourth, recent studies have shown that it is possible to activate somatotopic S1 hand representations through touch observation^69^ or attending to individual fingers^70^. As such, it is possible that simply observing others’ fingers being touched or attending to others’ finger movements may help to preserve somatotopic representations.

Our finding of preserved S1 somatotopy is seemingly inconsistent with the wealth of evidence showing cortical reorganisation in S1 following SCI.^5–8^ In these studies experimenters typically indirectly probe the deprived S1 hand cortex via stimulation of cortically adjacent body parts. Human fMRI studies similarly probed the intact and cortically neighbouring body parts and suggested that their representations shift to the deprived S1 cortex, though results are mixed.^9–14^ Further supporting research comes from transcranial magnetic stimulation (TMS) studies that inject current in localised areas of M1 to induce a peripheral muscle response. These studies demonstrated that the representations of less impaired muscles shift and expand following a complete or incomplete SCI, whereas representations of more impaired muscles retract or are absent.^14,71–75^ However, our fMRI results showed that SCI patients had a similar level of finger-related movement activity in M1 as controls, and preserved somatotopic hand representations in S1. It is possible that reorganisation and preservation of the original function could co-occur within cortical areas. Indeed, non-human primates demonstrated that remapping observed in S1 actually reflects reorganisation in subcortical areas of the somatosensory pathway, principally the brainstem.^6,76^ As such, the deprived S1 area receives reorganised somatosensory inputs upon tactile stimulation of neighbouring intact body parts. This would simultaneously allow the original S1 representation of the deprived body part to be preserved, as observed in our results when we directly probed the deprived S1 hand area through attempted finger movements.

Together, our findings indicate that in the first years after a tetraplegic SCI, the somatotopic S1 hand representation is preserved even in the absence of retained sensory function, motor function, and spared spinal tissue bridges. These preserved S1 finger maps could be exploited in a functionally meaningful manner by rehabilitation approaches that aim to establish new functional connections between the brain and the hand after an SCI, e.g. through neuroprosthetic limbs or advanced exoskeletons that are directly controlled by the brain.^23,77–79^

## Abbreviations

AIS: American Spinal Injury Association impairment scale
BET: brain extraction tool
BF: Bayes factor
FEAT: FMRI Expert Analysis Tool
FLIRT: FMRIB’s Linear Image Registration Tool
fMRI: functional MRI
FSL: FMRIB Software Library
FWHM: Full Width between Half Maximum
GLM: General Linear Model
GRASSP: Graded Redefined Assessment of Strength, Sensibility and Prehension
HRF: Haemodynamic Response Function
ISNCSCI: International Standards for Neurological Classification of Spinal Cord Injury
M1: primary motor cortex
MCFLIRT: Motion Correction using FMRIB’s Linear Image Registration Tool
S1: primary somatosensory cortex
SCI: spinal cord injury
TE: echo time
TMS: Transcranial magnetic stimulation
TR: repetition time

## Acknowledgements

We thank our participants for taking part in the study. We thank Lydia Kämpf, Nicolin Gauler, and Silvia Hofer for their assistance with data collection, Daan Wesselink for help with data analysis, and Tamar Makin for comments on the manuscript.

