## Supplementary Table 1 for "Finger somatotopy is preserved after tetraplegia but deteriorates over time"

| | Elbow flexion | Shoulder (deltoideus) | Wrist extension | Elbow extension | Fingers 2-5 extension | Thumb opposition | Thumb flexion | Middle finger flexion | Little finger abduction | Index finger abduction |
|---|---|---|---|---|---|---|---|---|---|---|
| **S01** | 5 | 4 | 0 | 0 | 0 | 0 | 0 | 0 | 0 | 0 |
| **S02** | 5 | 5 | 4 | 2 | 0 | 0 | 0 | 0 | 0 | 0 |
| **S03** | 4 | 4 | 4 | 3 | 1 | 0 | 0 | 0 | 0 | 0 |
| **S04** | 5 | 5 | 5 | 4 | 0 | 0 | 0 | 0 | 0 | 0 |
| **S05** | 4 | 1 | 3 | 4 | 3 | 3 | 1 | 1 | 1 | 2 |
| **S06** | 5 | 5 | 5 | 5 | 1 | 1 | 4 | 3 | 0 | 0 |
| **S07** | 5 | 5 | 5 | 5 | 1 | 1 | 1 | 0 | 1 | 1 |
| **S08** | 5 | 4 | 5 | 4 | 4 | 5 | 5 | 4 | 4 | 1 |
| **S09** | 5 | 4 | 4 | 4 | 4 | 4 | 5 | 5 | 4 | 4 |
| **S10** | 5 | 5 | 4 | 4 | 3 | 4 | 4 | 4 | 4 | 4 |
| **S11** | 5 | 5 | 4 | 2 | 5 | 4 | 5 | 5 | 1 | 1 |
| **S12** | 5 | 5 | 5 | 5 | 4 | 1 | 4 | 1 | 1 | 1 |
| **S13** | 5 | 4 | 4 | 4 | 5 | 4 | 5 | 3 | 4 | 4 |
| **S14** | 5 | 5 | 5 | 5 | 5 | 5 | 5 | 4 | 4 | 4 |

**Supplementary table 1: GRASSP motor sub-scores.** Each muscle was tested with resistance through its full range of motion and given a muscle grade between 0 and 5: 0 = flaccid motion, 1 = flicker motion, 2 = full range of motion with gravity eliminated, 3 = full range of motion against gravity, 4 = full range of motion with moderate resistance, 5 = full range of motion with maximal resistance.[1]
