## Supplementary Fig. 1 for "Finger somatotopy is preserved after tetraplegia but deteriorates over time"

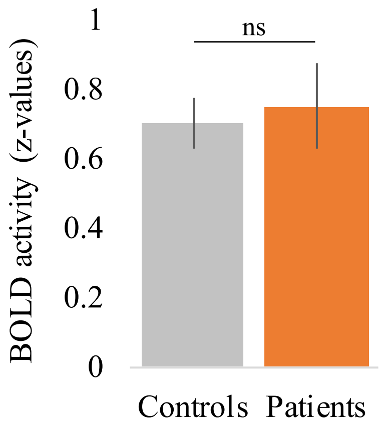


**Supplementary Figure 1:** We averaged the finger movement BOLD responses versus baseline across all fingers within the contralateral M1 hand ROI. Overall, all patients were able to engage their M1 hand area by moving individual fingers ($t_{(13)} = 6.16$, $p < 0.001$; $BF_{10} = 735$), as did controls ($t_{(17)} = 9.45$, $p < 0.001$; $BF_{10} = 3.82e+5$). Furthermore, patients' task-related activity was not significantly different from controls ($t_{(30)} = -0.34$, $p = 0.73$; $BF_{10} = 0.35$), with the BF showing anecdotal evidence in favour of the null hypothesis.
